# Whole genome assembly and annotation of the acorn weevil, *Curculio nanulus* (Coleoptera: Curculionidae)

**DOI:** 10.1101/2025.07.18.665623

**Authors:** Daniel D. Davis, Michael A. Charles, Duane D. McKenna, Paul B. Frandsen

## Abstract

The acorn weevil *Curculio nanulus* (Coleoptera: Curculionidae) is a seed predator that lays its eggs inside developing acorns and hickory nuts in the western United States. The female weevil uses her elongated rostrum to excavate a hole into the seed, creating a protected site for oviposition. Natural history traits among *Curculio* species—such as host-specificity and variation in larval diapause—suggest a dynamic evolutionary relationship with their host plants. These traits are best studied through a comparative genomic framework, but such analyses cannot currently be undertaken due to the lack of whole genome assemblies for *Curculio* species. To address this gap, we generated a whole genome assembly for *C. nanulus* using PacBio HiFi sequencing. The resulting assembly is ∼1.5 Gbp in length, with high contiguity (contig N50 = 7.7 Mbp) and gene completeness (USCO score: 97.46%). To enable comparative analysis, we also assembled the genome of the pecan weevil, *Curculio caryae*, using publicly available PacBio HiFi reads. For both species, we annotated repetitive elements and protein-coding genes, and compared these features with those of other weevil genomes. Our results reveal a marked expansion of repetitive elements within *Curculio* and its close relatives. These genomic resources provide a foundation for investigating seed predation, co-speciation, and host-parasite evolutionary dynamics in *Curculio* and related taxa, as well as their impacts on forest ecology.

## Introduction

Nut and acorn weevils of the genus *Curculio* (Coleoptera: Curculionidae) comprise a diverse lineage of seed predators with a distinctive life history. Females use their elongated rostrum to bore into developing hard-shelled seeds—typically those of oak, hazel, hickory, or chestnut—to deposit their eggs within the seed’s interior (Matsumura et al. 2021). This protected, nutrient-rich environment promotes larval development, but the presence of larvae simultaneously compromises the host seed’s viability, often preventing germination (Higaki 2016).

In response to such predation pressure, many oak species have evolved masting behavior—synchronized cycles of high and low seed production across populations (Kelly 1994). During mast years, trees produce a surplus of acorns, exceeding the consumption capacity of local seed predators like *Curculio*. In non-mast years, the reduced seed output curtails weevil reproduction, potentially lowering population densities (Higaki 2016). This cyclical mismatch between resource availability and consumer abundance represents an ecological and evolutionary strategy to limit seed predation.

*Curculio* species, in turn, have evolved counter-adaptations. After larvae exit their host seeds, they enter a period of diapause within the soil, which may last for one or more years. This dormancy allows populations to persist through low-resource periods and potentially synchronize emergence with future mast events. The duration and dynamics of diapause vary widely both among and within *Curculio* species (Higaki 2016; Espelta et al. 2017; Menu and Desouhant 2002), suggesting ongoing evolutionary adaptation to host reproductive cycles (Maeto and Ozaki 2003).

This co-evolutionary relationship between host and seed predator has likely shaped key traits such as host specificity, life cycle timing, and ecological resilience. Despite their ecological significance, the genetic basis of these traits in *Curculio* remains poorly understood. The lack of whole genome assemblies for *Curculio* species has prevented investigations into the molecular underpinnings of diapause regulation, host-plant interactions, and adaptive evolution.

To address this, we sequenced and assembled the genome of *Curculio nanulus*, a western U.S. acorn weevil, using PacBio HiFi technology. To enable comparative analyses, we also assembled the genome of the pecan weevil, *Curculio caryae,* using publicly available PacBio HiFi reads generated by the United States Department of Agriculture (NCBI SRA: SRR18245025). These assemblies serve as a foundation for future evolutionary and ecological studies in *Curculio* and other seed-parasitic insects.

## Methods

### DNA Extraction and Sequencing

We collected a larval specimen of *Curculio nanulus* from an acorn found underneath an oak tree on the Brigham Young University campus in Provo, UT, USA, in September 2022. We extracted high molecular weight (HMW) genomic DNA from the specimen using the Qiagen genomic-tip DNA extraction kit. We sheared purified DNA to 18 kbp with a Diagenode Megaruptor and used a BluePippin system (Sage Science, Beverly, MA, USA) to collect fractions containing >15 kbp fragments for library preparation. We prepared genomic DNA libraries using the SMRTbell Express Template Prep Kit 2.0 and associated protocol (PacBio, Menlo Park, CA, USA) and sequenced the library on a single PacBio Revio flow cell at the BYU DNA Sequencing Center. Additionally, we downloaded publicly available PacBio HiFi sequencing reads from the NCBI SRA for the pecan weevil, *Curculio caryae* (SRR18245025), to support comparative analyses.

### Assembly and Refinement

We assembled the PacBio HiFi reads for each species using hifiasm v0.16.1-r375 (Cheng et al. 2021). We subsequently used purge_dups v1.2.5 to remove duplicated haplotigs (Guan et al. 2020). Purge_dups uses contig similarity and depth of coverage to assess whether a contig represents haplotypic variation rather than a true duplication. Contigs identified as haplotypic duplicates were excluded from the assembly.

We identified and removed contaminant contigs using BlobTools v1.1.1 (Laetsch and Blaxter 2017; Laetsch et al. 2017) based on closest taxonomic match from a blastn v2.12.0 search against the NCBI nt database, GC proportion, and sequencing coverage.

We located, circularized, and annotated the mitochondrial genomes for each species using MitoHiFi (Uliano-Silva et al. 2023) with the maize weevil, *Sitophilus zeamais*, (NC_030764.1, 18105 bp, 37 genes) (Ojo et al. 2016) mitochondrial genome as a reference. Circular representations of the contigs were visualized with OrganellarGenomeDraw v1.3.1 (Greiner, Lehwark, and Bock 2019).

### Identification

Following whole genome sequencing, we discovered that the sequenced larval specimen could not be identified to the species-level using a morphological key because no such key exists for the *Curculio* larvae of the western United States. To aid in our identification, we collected larval specimens from the same locality and reared the larvae to adulthood in a moist coconut coir substrate kept between 65 and 72°F. We identified adult specimens to species using the key from Gibson (1969) and imaged the specimens with an Olympus DP75 camera mounted on a SZX-ILLB100 microscope (Fig. S1). We generated COI-5P barcodes for two identified adult and two larval specimens using the primers LCO1490-L and HCO2198-L (Nelson, Wallman, and Dowton 2007), to aid in confirming the identification of the sequenced specimen.

To determine whether the species from which we sequenced the whole genome was the same species as those we reared, we extracted the COI gene from the mitochondrial genome assembly and aligned it with the COI-5P barcodes from the identified specimens using MUSCLE as implemented in MEGA v11.0.13 (Tamura, Stecher, and Kumar 2021). MEGA v11.0.13 was used both to calculate pairwise distances and to construct the maximum likelihood tree using default parameters.

### Quality Control

For each purged and decontaminated genome assembly, we used assembly_stats.py (Trizna 2020) to measure assembly statistics including N50, L50, genome size, number of contigs, and GC content. The average sequencing coverage was calculated for each assembly by dividing the assembly size by the total read lengths of the respective sequencing runs.

We used compleasm to estimate gene completeness, by estimating the number of universal single-copy orthologs (USCOs) that each genome shares with the endopterygota_odb10 ortholog set (Huang and Li 2023).

### Genome Annotation

We identified repetitive regions in each genome *ab initio* with RepeatModeler v2.0.5 and subsequently soft-masked repetitive regions using RepeatMasker v4.1.5 with the default parameters (Flynn et al. 2020). We conducted further analysis of repetitive elements with Earl Grey v4.4.0 (Baril, Galbraith, and Hayward 2024).

We completed structural annotations on the soft-masked assemblies using the GALBA v1.0.11 pipeline (Brůna et al. 2023; H. Li 2023; Buchfink, Xie, and Huson 2015; Stanke et al. 2006; Hoff and Stanke 2019) with the protein annotation from the red palm weevil *Rhynchophorus ferrugineus* as a reference (Sudalaimuthuasari et al. 2024). Completeness of each annotation was evaluated with compleasm v0.2.5 (Huang and Li 2023) in protein mode with the endopterygota_odb10 ortholog set. Functional annotations were completed using Blast2GO v1.5.1 (Conesa and Götz 2008).

### Comparison of repetitive elements

We used Earl Grey v4.4.0 (Baril, Galbraith, and Hayward 2024) to annotate the repetitive elements of six additional curculionid genomes and one brentid genome available from GenBank (Table 1). We arranged visualizations of the repetitive elements on a phylogeny constructed using the protein sequences from compleasm annotations. To construct the phylogeny, we aligned the amino acid sequences from each single-copy compleasm locus with MAFFT v7.526. We scored the alignments with Aliscore v02.2 and trimmed them with ALICUT V2.31. We estimated the final phylogenetic tree using two methods: first, by concatenating the aligned sequences and conducting a partitioned model search followed by maximum likelihood construction with IQtree v2.1.3; and second, by constructing a gene tree for each locus with IQtree v2.1.3 followed by species tree estimation with Astral III v5.7.1, using the gene trees as input. The phylogeny was rooted using *Cylas formicarius* (Brentidae: Cyladinae) and then visualized using FigTree v1.4.4 (http://tree.bio.ed.ac.uk/software/figtree/). We arranged plots showing the change in the repetitive elements over time for each species on the phylogeny.

**Table 1.** Genomes used in comparison of repetitive elements.

| **Table 1** Genomes used in comparison of repetitive elements. |  |  |
| --- | --- | --- |
| **Family** | **Species** | **GenBank Accession** |
| Curculionidae | *Anthonomus grandis grandis* | GCA_022605725.3 |
|  | *Dendroctonus valens* | GCA_024550625.1 |
|  | *Curculio caryae* | Assembled from SRR18245025 |
|  | *Curculio nanulus* | (assembled for this study) |
|  | *Kuschelorrhynchus macadamiae* | GCA_030620095.1 |
|  | *Listronotus oregonensis* | GCA_019359885.1 |
|  | *Rhynchophorus ferrugineus* | GCA_030347505.1 |
|  | *Sitophilus oryzae* | GCA_002938485.2 |
| Brentidae | *Cylas formicarius* (outgroup) | GCA_029955315.1 |

### Comparison to available Curculionidae genomes

There are 38 genome assemblies representing 24 species belonging to the weevil family Curculionidae in GenBank. We evaluated all 38 genomes for contiguity using assembly_stats.py (Trizna 2020) and completeness using compleasm with the endopterygota_odb10 ortholog set. For downstream analyses, we included the best genome for each species based on the compleasm completeness statistic. The curculionid genomes that we used are available at https://www.ncbi.nlm.nih.gov/datasets/genome/?taxon=7042.

## Results

### Species verification

The COI sequence from the *Curculio nanulus* whole genome assembly differed by no more than 0.3% from verified *C. nanulus* specimens (Table S1). Phylogenetic analysis also placed our sequenced specimen (GenBank: JBEWYK010000000) firmly within the *C. nanulus* clade (Fig. S2), strongly supporting its species identity.

### Genome sequencing and assembly

Sequencing of *C. nanulus* generated 81.8 Gbp of data across 5,212,942 PacBio HiFi reads. The initial assembly (Table 2) totaled 1.6 Gbp across 898 contigs, with 51x sequencing coverage, a contig L50 of 57 contigs, and a contig N50 of 7.19 Mbp. Following refinement via purge_dups and BlobTools, the assembly was reduced to 1.5 Gbp across 652 contigs, improving contiguity (N50 = 7.71 Mbp; L50 = 51 contigs) and sequencing coverage (54x).

**Table 2.** Contiguity and completeness statistics for the *Curculio nanulus* initial, intermediate and final genome assemblies as well as completeness for gene annotations. Sequencing coverage was calculated by dividing the number of base pairs in the raw sequencing reads by the assembly size. Compleasm was run with the endopterygota_odb10 database.

|  | *Curculio nanulus* |  |  |
| --- | --- | --- | --- |
| **Sequencing reads (bp)** | 81,847,322,991 |  |  |
| **Reads count** | 5,212,942 |  |  |
| **Mean read length** | 15,700.8 |  |  |
| **Read N50** | 15,830 |  |  |
|  | Initial Assembly | Post-PurgeDups | Decontaminated with Blobtools |
| **Contig count** | 898 | 654 | 652 |
| **L50** | 57 | 51 | 51 |
| **N50** | 7,189,678 | 7,709,206 | 7,709,206 |
| **% GC content** | 35.059 | 35.068 | 35.073 |
| **Shortest contig (bp)** | 5,225 | 5,225 | 5,225 |
| **Mean contig (bp)** | 1,785,266 | 2,311,745.6 | 2,316,901.6 |
| **Median contig (bp)** | 355,929.5 | 549,145.5 | 549,145.5 |
| **Longest contig (bp)** | 24,979,329 | 24,979,329 | 24,979,329 |
| **Total length (bp)** | 1,603,168,799 | 1,511,881,652 | 1,510,619,856 |
| **sequencing coverage** | 51x | 54x | 54x |
| **USCOs (compleasm)** |  |  |  |
| Complete (%) | 99.01 | 98.97 | 98.97 |
| Single copy (%) | 95.90 | 97.46 | 97.46 |
| Duplicated (%) | 3.11 | 1.51 | 1.51 |
| Fragmented (%) | 0.24 | 0.24 | 0.24 |
| Missing (%) | 0.75 | 0.80 | 0.80 |
| **Annotation** |  |  |  |
| Putative genes | 19,235 |  |  |
| Complete (%) | 95.34 |  |  |
| Single copy (%) | 79.10 |  |  |
| Duplicated (%) | 16.24 |  |  |
| Fragmented (%) | 2.54 |  |  |
| Missing (%) | 2.12 |  |  |

After refinement, the *C. nanulus* assembly contained 98.97% complete universal single copy orthologs (USCOs). Complete and single-copy USCOs increased from 95.90% to 97.46%, while duplicated USCOs decreased from 3.11% to 1.51%, indicating improved assembly quality and reduced haplotypic duplication, with only a minimal (∼0.04%) decrease in total completeness.

The mitochondrial genome of *C. nanulus* was 18,823 bp in size (Fig. 3), encoding 37 genes, the same number present in the reference and *C. caryae* (20,075 bp). Gene order was generally conserved between *C. nanulus* and *C. caryae*, with only minor differences in the positions of tRNAs relative to the reference.

The *C. nanulus* assembly is highly contiguous, and contiguity improved with post-assembly refinement using purge_dups (Guan et al. 2020). Notably, despite their close relationship, the refined *C. nanulus* assembly was ∼1.51 Gbp—approximately 634 Mbp smaller than the *C. caryae* assembly (∼2.14 Gbp; Table S2).

Further results for the *C. caryae* assembly can be found in the supplemental material.

### Repetitive elements and genome size

The *C. nanulus* assembly was 79.11% repetitive (1,195.1 Mbp), whereas the *C. caryae* assembly was 84.16% repetitive (1,805.1 Mbp) (Fig. 1). The sizes of non-repetitive regions were comparable: 315.5 Mbp in *C. nanulus* and 339.8 Mbp in *C. caryae* (Fig. 2B). Therefore, the 634 Mbp difference in total genome size is attributable primarily to greater expansion in the repetitive content of the *C. caryae* genome, especially unclassified repeats, DNA transposons, LINEs, and LTR retrotransposons. In both *Curculio* genomes, the majority of the repetitive elements were unclassified: 34.6% of the *C. nanulus* genome and 30.7% of the *C. caryae* genome.

**Figure 1.**
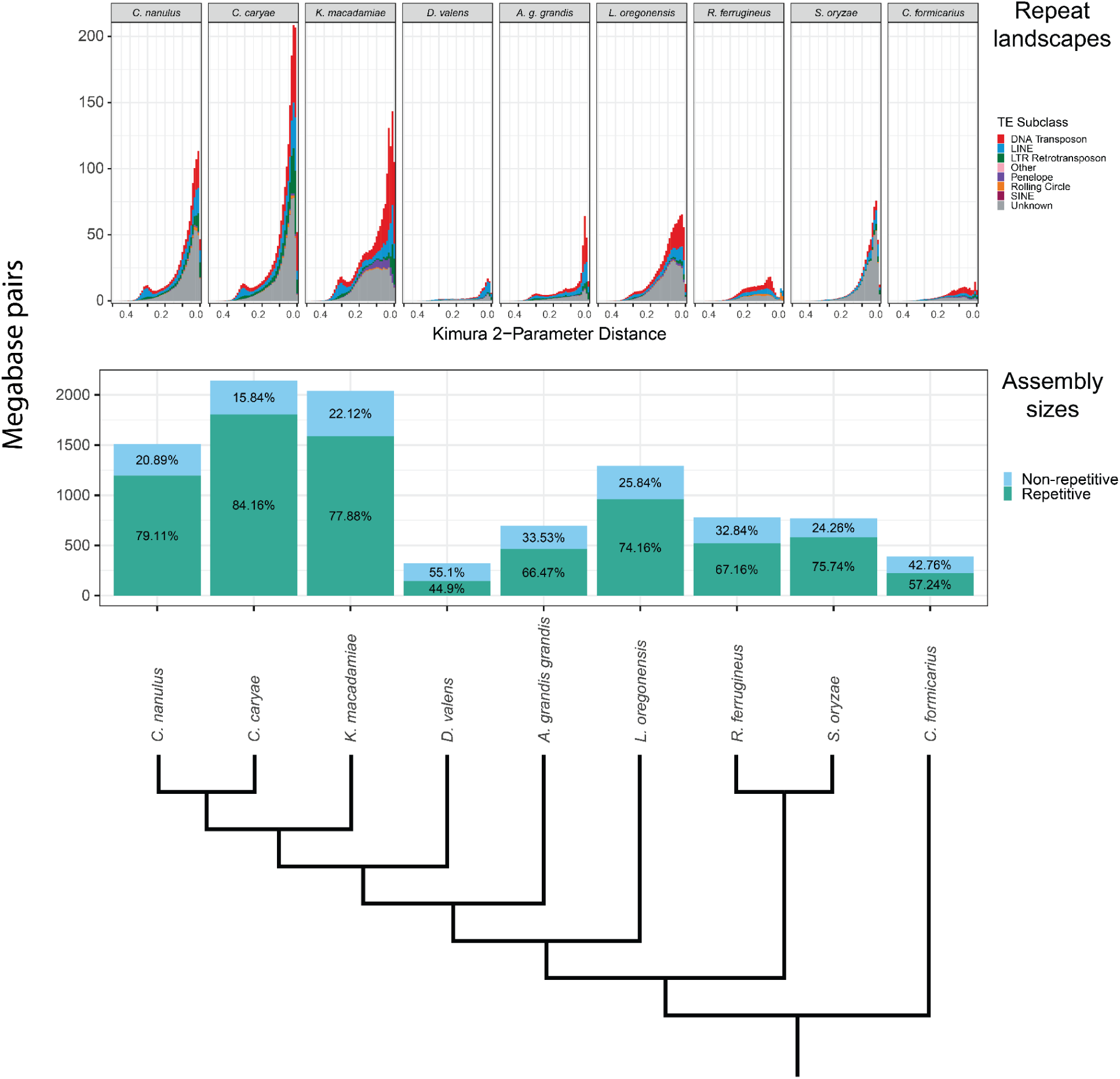
Repetitive elements identified in the genomes assemblies of eight curculionid and one brentid species. The species are ordered according to the cladogram shown. The top plot shows changes in total repetitive elements across time. The second plot compares genome assembly sizes broken into repetitive and non-repetitive DNA.

**Figure 2.**
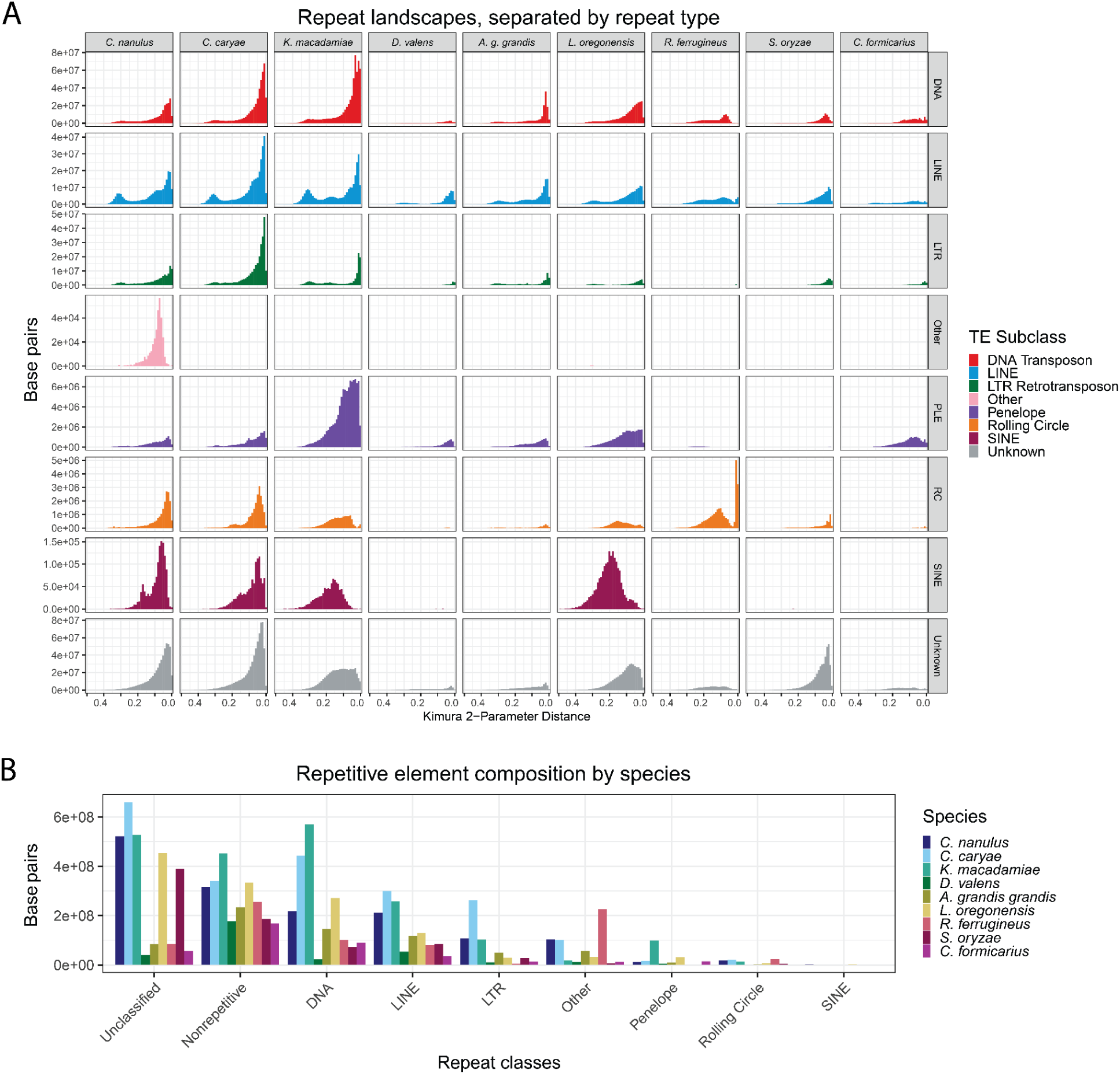
Repetitive elements of eight curculionid and one brentid species. **A)** Repetitive elements across time split into classes. **B)** Comparison of total base pairs of major repeat classes broken down by species. “Other” includes simple repeats, microsatellites, and repetitive RNA.

The maximum likelihood tree constructed from concatenated sequences and the ASTRAL species tree had identical topologies. Mapping repeat landscapes onto this phylogeny (Fig. 1) showed a shared recent expansion of repetitive elements in the clade containing *C. nanulus*, *C. caryae*, and *Kuschelorhynchus macadamiae*. This expansion continued within *Curculio,* though certain repeat types (e.g., unclassified elements, SINEs, and rolling-circle elements) plateaued in *K. macadamiae* (Fig. 2A). In contrast, *K. macadamiae* exhibited substantial growth of Penelope elements, which now occupy 6-9 times more sequence than in *Curculio,* yet still comprise just 4.9% of the genome.

Among *Curculio*, *C. caryae* exhibited greater repetitive element expansion than *C. nanulus*, explaining its larger genome size. Except for *Dendroctonus valens*, all analyzed weevil genomes contained at least 57% repetitive content (Fig. 1). Independent repeat expansions were evident in the lineages leading to *A. grandis grandis*, *L. oregonensis*, *R. ferrugineus,* and *S. oryzae,* distinct from the *Curculio/Kuschelorhynchus* expansion.

In *C. nanulus*, unclassified elements comprised 43.7% of repetitive elements (34.6% of the genome), while in *C. caryae*, they accounted for 36.6% of repetitive elements (30.7% of the genome). This pattern aligns with findings from other weevils (e.g., Sylvester et al. 2024) and other non-model insects, where an average of 40.5% of repeats remain undescribed outside of *Drosophila* (Sproul et al. 2023).

### Gene annotation

The *C. nanulus* genome annotation yielded 19,235 genes using the GALBA pipeline, with a single-copy USCO completeness of 79.10% and duplicated completeness of 16.24%, for a total completeness of 95.34% (Table 2). Similarly, in *C. caryae*, 20,473 genes were annotated, with 78.39% single-copy completeness, and 15.91% duplicated completeness, for a total completeness of 94.84% (Table S2). For comparison, the reference annotation used in the GALBA pipeline, *Rhynchophorus ferrugineus* (PRJNA950221), included 29,666 genes with 98.26% total USCO completeness (77.64% single-copy and 20.62% duplicated).

Blast2GO assigned Gene Ontology (GO) terms to 85% of predicted *C. nanulus* genes, with 72% receiving functional annotations. For *C. caryae,* 84% of genes were assigned GO terms, and 69% were functionally annotated.

### Genome Comparison

The *C. nanulus* assembly ranked among the highest-quality of the 24 publicly available curculionid genomes. It was amongst the four highest contig N50s and nine highest USCO completeness scores (Table S3).

## Discussion

### Genomic resources for studying co-evolution

Species evolve within ecological networks shaped by complex webs of interspecies interactions. For seed-parasitic insects like *Curculio*, evolutionary trajectories are intimately tied to their host plants. The newly assembled genome of *C. nanulus* provides an essential resource for studying co-evolutionary processes at the genomic level.

*Curculio* species can profoundly influence forest biodiversity by reducing the reproductive success of dominant tree species, potentially benefiting understory or competing plants (Q.-Y. Li et al. 2021). The ecological consequences of this predation depend on the host species targeted and the broader forest context. High-quality genome assemblies offer a means to investigate the genomic basis of plant-feeding habits in seed predators, and how the actions of such herbivores shape plant communities.

This assembly also enables investigation into the genomic basis of diapause, a key adaptation in *Curculio*. Diapause synchronizes weevil emergence with host seed availability, enabling persistence through masting cycles. Genes regulating circadian rhythms have been implicated in diapause control in arthropods such as *Drosophila* (Yamada and Yamamoto 2011) and *Daphnia* (Schwarzenberger, Chen, and Weiss 2020). Circadian clock genes such as these are likely candidates for similar diapause-regulating functions in *Curculio*. Understanding how these genes evolve in response to host phenology may reveal how quickly *Curculio* can shift hosts in response to environmental change or host decline. This, in turn, will improve predictions of co-extinction risk and resilience in host-parasite systems.

### Repetitive Elements

Repetitive elements are a dominant feature of eukaryotic genomes and play a central role in genome evolution, regulation, and architecture (Bourque et al. 2018; Gilbert, Peccoud, and Cordaux 2021). For instance, repetitive elements have been found to influence 3-dimensional genomic architecture in ground beetles, as well as strongly reflect species boundaries (Sproul, Barton, and Maddison 2020). Our analysis shows that repeat expansion is a defining feature of *Curculio* genome evolution. The substantial repeat content—particularly unclassified elements—suggests that novel or rapidly evolving repeat families contribute significantly to genome size differences within the genus.

Repeat expansion in *Curculio* and its close relatives appears to have occurred independently of other curculionid lineages, indicating a lineage-specific dynamic. Notably, the large difference in genome size between *C. nanulus* and *C. caryae* can be attributed almost entirely to differences in repeat content, especially unclassified elements, LINEs, DNA transposons, and LTR retrotransposons. These genome assemblies and annotations are valuable resources for characterizing novel repeats in non-model insects and understanding how repeat dynamics influence genome evolution, species divergence, and adaptation in the species rich family Curculionidae.

## Data Availability

This Whole Genome Shotgun project for *Curculio nanulus* has been deposited at DDBJ/ENA/GenBank under the accession JBEWYK000000000. The version described in this paper is version JBEWYK010000000. The PacBio HiFi reads are available under BioProject PRJNA1129540.

Reviewer Link: https://dataview.ncbi.nlm.nih.gov/object/PRJNA1129540?reviewer=vnloe7khr72o2f09rij27sbm7f

Annotations and intermediate assemblies are available at https://byu.box.com/s/yzplmri8g6fwm6v5zhqrvpjd84azz21h (will be uploaded to Figshare if manuscript is accepted)

The PacBio HiFi reads generated by the Ag100Pest Initiative for *Curculio caryae* are available in sequencing run SRR18245025.

## Acknowledgements

We would like to thank the Ag100Pest Initiative for generating the sequencing reads for *Curculio caryae* (SRR18245025), the Brigham Young University DNA Sequencing Center for extracting and sequencing the DNA from the provided *Curculio nanulus* sample, and Riley Nelson for his photography training.

## Supplemental Material

### Results for *C. caryae* assembly

Publicly available PacBio HiFi reads for *Curculio caryae* totaled 92.7 Gbp across 10,074,085 reads. The initial assembly (Table S2) consisted of 2.2 Gbp across 1,146 contigs, with 42x coverage, a contig L50 of 131, and a contig N50 of 5.02 Mbp. After refinement, the assembly length was reduced to 2.1 Gbp across 826 contigs, with slightly improved metrics: 43x coverage, contig L50 of 125, and contig N50 of 5.12 Mbp.

Completeness metrics for the *C. caryae* assembly remained high after refinement. While the total USCO completeness score decreased from 98.82% to 98.59%, complete and single-copy USCOs rose from 97.41% to 97.79%, and duplicated USCOs dropped from 1.41% to 0.80%. Fragmented and missing genes remained unchanged, indicating minimal compromise to completeness while removing redundancy and contaminants.

**Figure S1.**
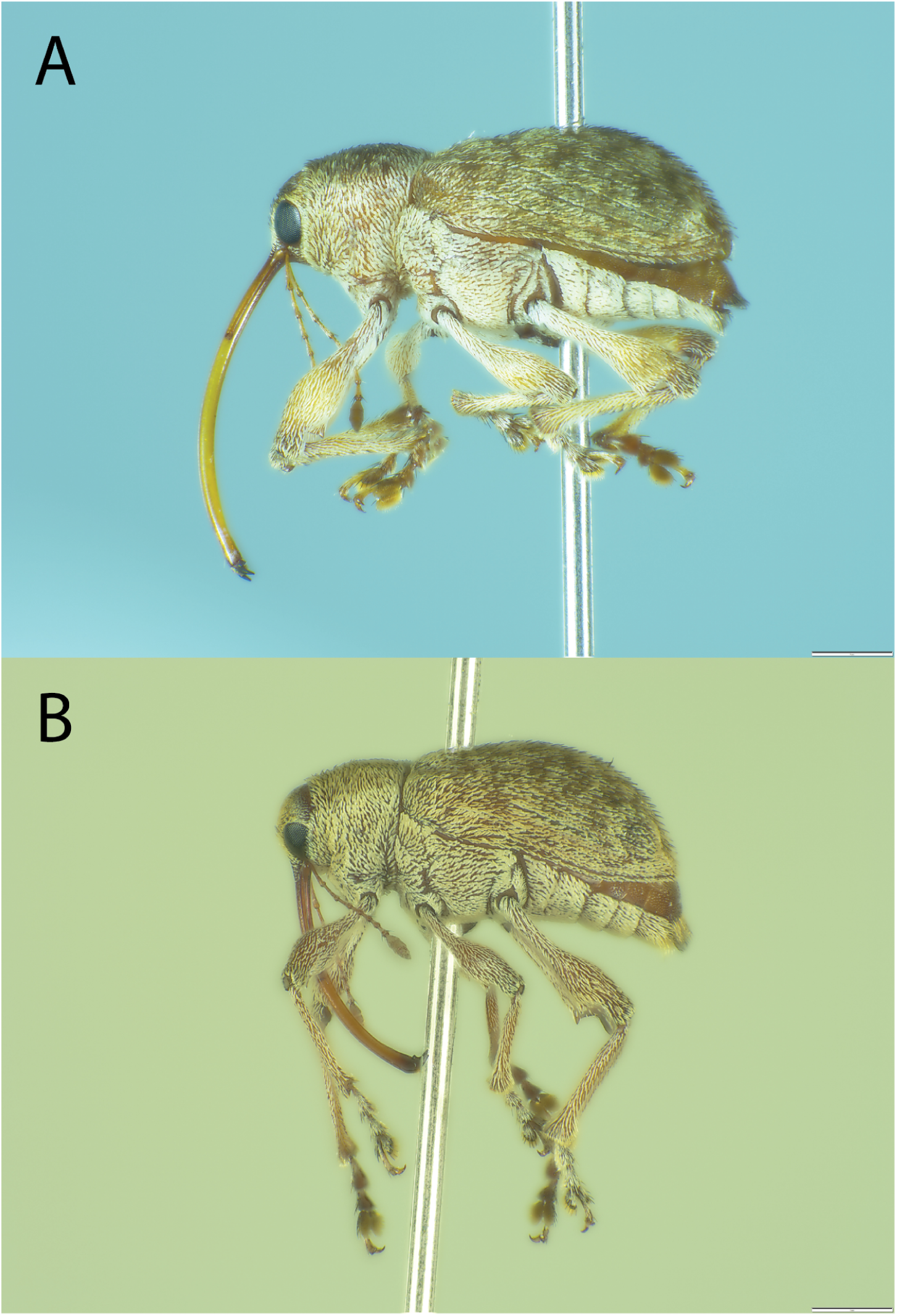
Photographs of *Curculio nanulus.* **A)** Lateral view of adult (specimen V0017). **B)** Lateral view of adult (specimen V0024).

**Figure S2.**
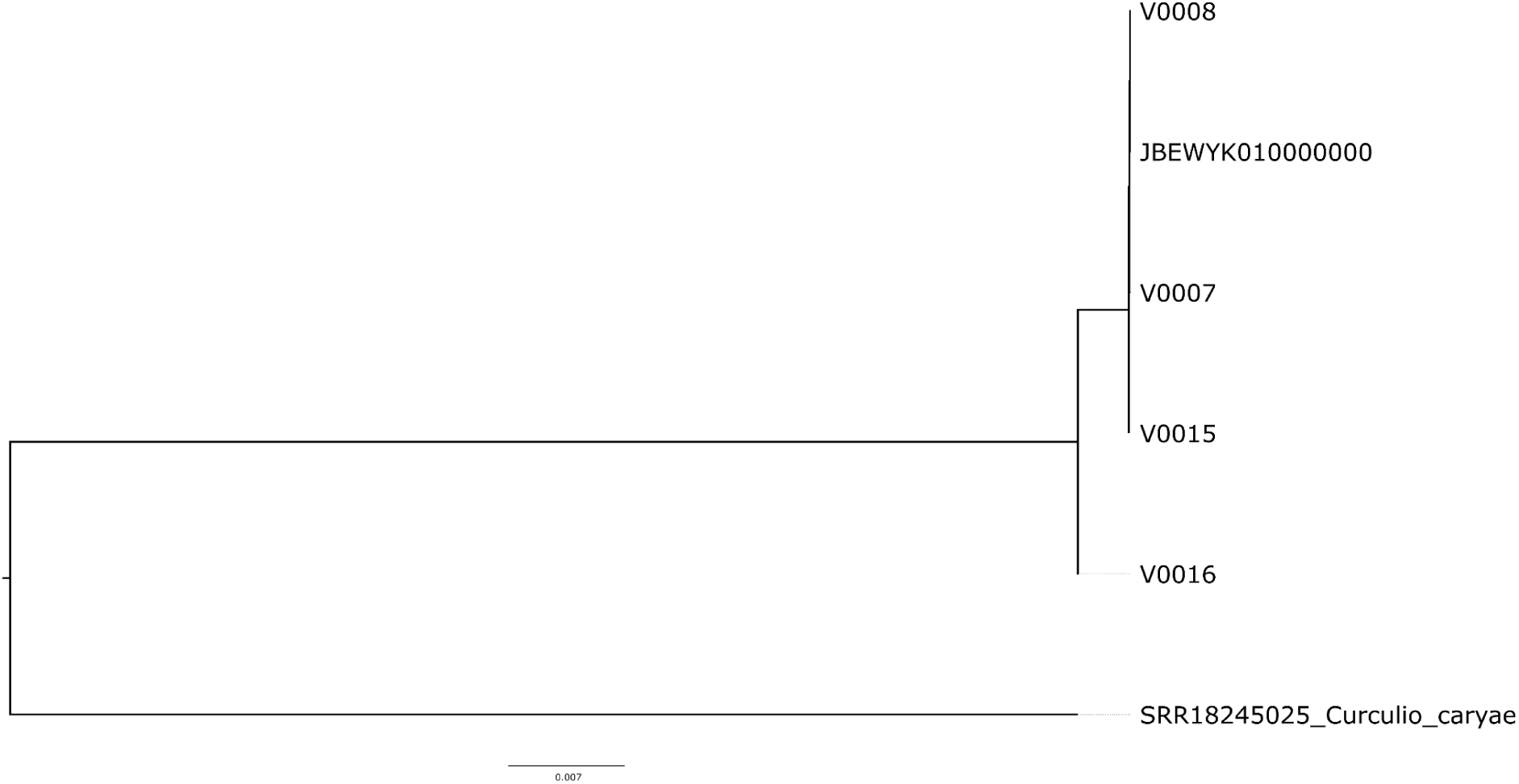
Phylogeny of *Curculio nanulus* specimens constructed using COI-5P barcodes placing the genome specimen (JBEWYK010000000) within *C. nanulus.* V0007 and V0008 are larval specimens collected from the same tree concurrent with JBEWYK010000000. V0015 and V0016 were collected from a nearby oak tree, reared to adulthood, and identified to the species level.

**Figure S3.**
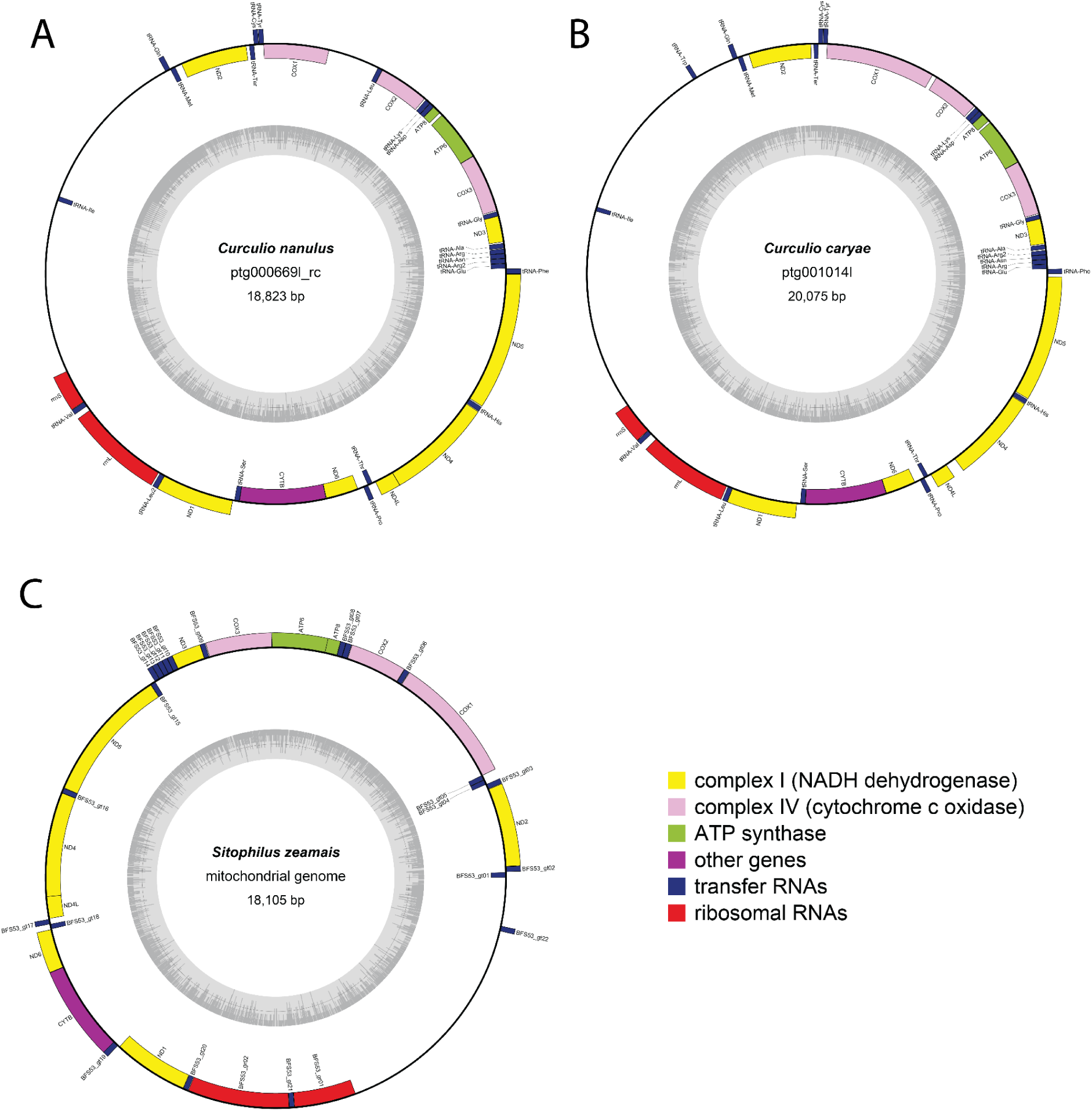
Visualisations of the mitochondrial genome assemblies generated with OrganellarGenomeDraw v1.3.1. A) *Curculio nanulus*; B) *C. caryae*; C) *Sitophilus zeamais*, the reference mitochondrial genome.

**Table S1.**
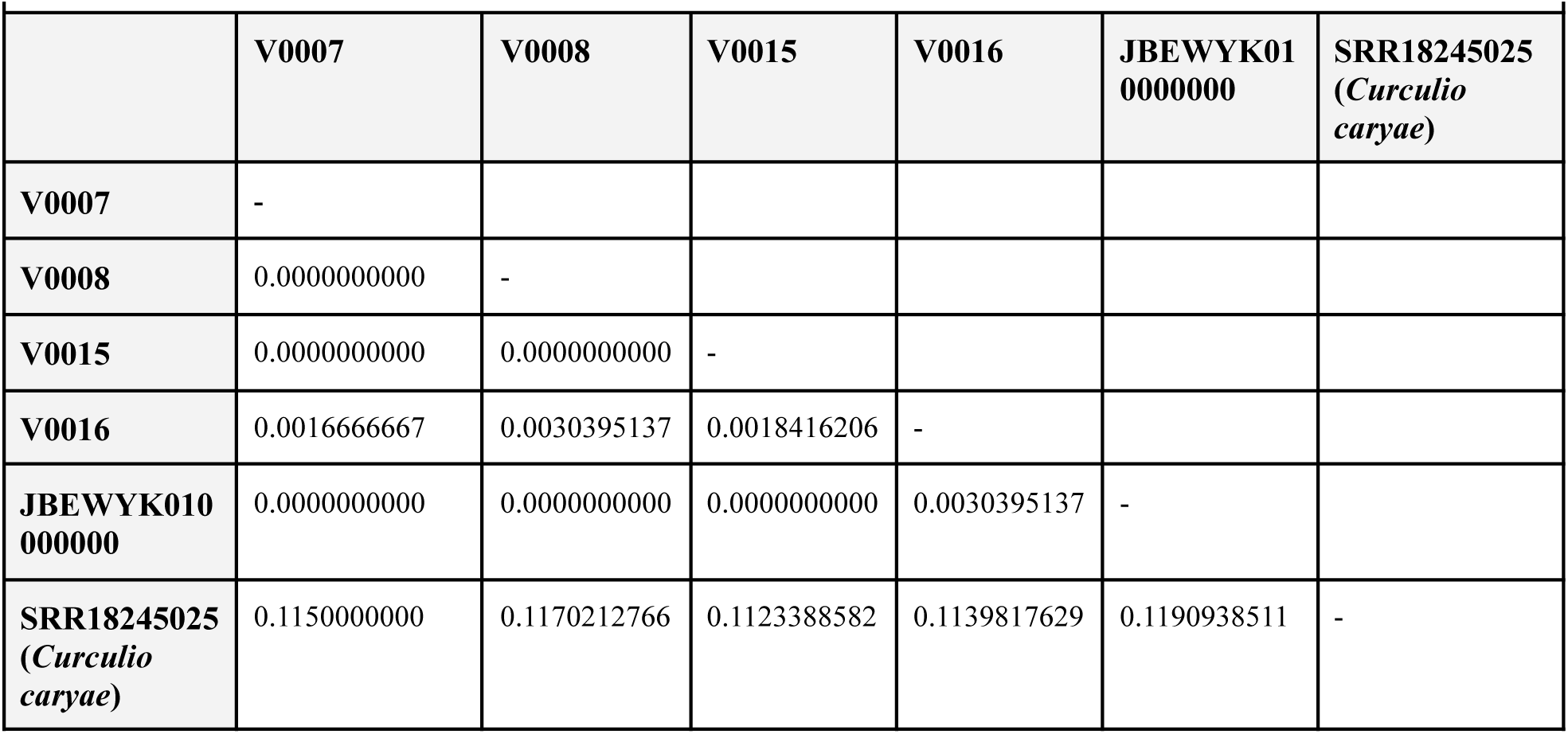
Pairwise distances between the COI-5P barcodes of identified *C. nanulus* specimens and the COI gene extracted from the likely *C. nanulus* genome.

**Table S2.** Contiguity and completeness statistics for the *Curculio caryae* initial, intermediate and final genome assemblies as well as completeness for gene annotations. Sequencing coverage was calculated by dividing the number of base pairs in the raw sequencing reads by the assembly size. Compleasm was run with the endopterygota_odb10 database.

|  | *Curculio caryae* |  |  |
| --- | --- | --- | --- |
| **Sequencing reads (bp)** | 92,686,342,389 |  |  |
| **Reads count** | 10,074,085 |  |  |
| **Mean read length** | 9,200.5 |  |  |
| **Read N50** | 10,381 |  |  |
|  | **Initial Assembly** | **Post-PurgeDups** | **Decontaminated with Blobtools** |
| **Contig count** | 1146 | 864 | 826 |
| **L50** | 131 | 128 | 125 |
| **N50** | 5,021,767 | 5,099,020 | 5,121,652 |
| **% GC content** | 35.4059 | 35.397 | 35.160 |
| **Shortest contig (bp)** | 5,207 | 5,207 | 5,339 |
| **Mean contig (bp)** | 1,927,898.5 | 2,521,315.3 | 2,596,665.7 |
| **Median contig (bp)** | 841,781 | 1,434,915 | 1,502,193 |
| **Longest contig (bp)** | 27,825,805 | 27,825,805 | 27,825,805 |
| **Total length (bp)** | 2,209,371,675 | 2,178,416,419 | 2,144,845,900 |
| **sequencing coverage** | 42x | 43x | 43x |
| **USCOs (compleasm)** |  |  |  |
| Complete (%) | 98.82 | 98.73 | 98.59 |
| Single copy (%) | 97.41 | 97.79 | 97.79 |
| Duplicated (%) | 1.41 | 0.94 | 0.80 |
| Fragmented (%) | 0.56 | 0.66 | 0.61 |
| Missing (%) | 0.61 | 0.61 | 0.80 |
| **Annotation** |  |  |  |
| Putative genes | 20,473 |  |  |
| Complete (%) | 94.84 |  |  |
| Single copy (%) | 78.39 |  |  |
| Duplicated (%) | 15.91 |  |  |
| Fragmented (%) | 2.87 |  |  |
| Missing (%) | 2.82 |  |  |

**Table S3.** Assembly statistics for each of the 24 species of weevil (Curculionidae) genomes represented in GenBank, as well as the two *Curculio* genomes assembled in this study (rows bolded and highlighted). Rows are ordered by Contig N50 measured in kilobase pairs.

| Assembly Accession | Organism Name | USCO Complete | Coverage | Contig L50 | Contig N50 (Kbp) | GC content | Contig count | Genome Size (Mbp) |
| --- | --- | --- | --- | --- | --- | --- | --- | --- |
| GCA_030347505.1 | *Rhynchophorus ferrugineus* | 99.25% | 100 | 13 | 20,155.5 | 33.671 | 744 | 778.73 |
| GCA_935413205.1 | *Polydrusus cervinus* | 99.34% | 38 | 12 | 19,113.4 | 32.769 | 75 | 713.36 |
| GCA_030068095.1 | *Anthonomus grandis thurberiae* | 99.01% | 11 | 25 | 9,069.1 | 32.276 | 754 | 738.35 |
| **JBEWYK000000000** | ***Curculio nanulus*** | **98.90%** | **54** | **51** | **7,709.2** | **35.073** | **652** | **1,510.62** |
| GCA_022605725.3 | *Anthonomus grandis grandis* | 99.01% | 21 | 31 | 6,866.0 | 32.130 | 568 | 697.43 |
| GCA_016097725.1 | *Ips typographus* | 99.20% | 211 | 11 | 6,654.0 | 35.207 | 272 | 236.82 |
| **SRR18245025** | ***Curculio caryae*** | **98.59%** | **43** | **125** | **5,121.7** | **35.160** | **826** | **2,144.85** |
| GCA_917834065.1 | *Ceutorhynchus assimilis* | 98.21% | 40 | 51 | 3,538.5 | 35.294 | 628 | 674.66 |
| GCA_949748235.1 | *Platypus cylindrus* | 96.99% | 181 | 18 | 2,446.3 | 31.055 | 128 | 147.46 |
| GCA_963576705.1 | *Polydrusus tereticollis* | 99.01% | 15 | 274 | 1,537.9 | 34.042 | 1827 | 1,392.42 |
| GCA_958502075.1 | *Orchestes rusci* | 99.11% | 34 | 144 | 1,248.7 | 35.592 | 933 | 623.88 |
| GCA_951812265.1 | *Taphrorychus bicolor* | 99.20% | 31 | 128 | 1,227.0 | 36.233 | 1,283 | 575.09 |
| GCA_030620095.1 | *Kuschelorrhynchus macadamiae* | 96.61% | 98 | 594 | 989.5 | 31.862 | 5,521 | 2,040.83 |
| GCA_024550625.1 | *Dendroctonus valens* | 96.47% | 806 | 82 | 985.5 | 36.833 | 1,143 | 320.96 |
| GCA_002938485.2 | *Sitophilus oryzae* | 98.73% | 101 | 257 | 723.2 | 32.632 | 5,424 | 757.90 |
| GCA_031761425.1 | *Cosmopolites sordidus* | 97.98% | 20 | 442 | 706.4 | 33.974 | 2,977 | 1,068.54 |
| GCA_019049505.1 | *Pachyrhynchus sulphureomaculatus* | 93.13% | 40 | 2,082 | 299.7 | 33.752 | 14,365 | 2,050.36 |
| GCA_018691245.2 | *Ips nitidus* | 96.23% | 94 | 285 | 175.9 | 35.173 | 33,250 | 231.34 |
| GCA_014170235.1 | *Listronotus bonariensis* | 94.35% | 38 | 2,726 | 120.3 | 31.308 | 22,298 | 1,112.42 |
| GCA_019359885.1 | *Listronotus oregonensis* | 87.62% | 67 | 6,281 | 53.1 | 30.955 | 44,498 | 1,292.90 |
| GCA_020466585.2 | *Dendroctonus ponderosae* | 97.93% | 747 | 2,136 | 26.1 | 35.803 | 20,279 | 213.86 |
| GCA_016904865.1 | *Pissodes strobi* | 80.79% | 70 | 21,576 | 24.9 | 35.390 | 144,278 | 1,831.48 |
| GCA_001012855.1 | *Hypothenemus hampei* | 97.70% | 100 | 18,102 | 1.9 | 32.255 | 219,563 | 130.40 |
| GCA_014849505.1 | *Elaeidobius kamerunicus* | 40.49% | 17 | 92,726 | 0.8 | 31.703 | 478,729 | 263.84 |

## References

1. Baril, Tobias, James Galbraith, and Alex Hayward. 2024. “Earl Grey: A Fully Automated User-Friendly Transposable Element Annotation and Analysis Pipeline.” Molecular Biology and Evolution 41 (4): msae068. 10.1093/molbev/msae068.

2. Bourque, Guillaume, Kathleen H. Burns, Mary Gehring, Vera Gorbunova, Andrei Seluanov, Molly Hammell, Michaël Imbeault, et al. 2018. “Ten Things You Should Know about Transposable Elements.” Genome Biology 19 (1): 199. 10.1186/s13059-018-1577-z.

3. Brůna, Tomáš, Heng Li, Joseph Guhlin, Daniel Honsel, Steffen Herbold, Mario Stanke, Natalia Nenasheva, Matthis Ebel, Lars Gabriel, and Katharina J. Hoff. 2023. “GALBA: Genome Annotation with Miniprot and AUGUSTUS.” bioRxiv. 10.1101/2023.04.10.536199.

4. Buchfink, Benjamin, Chao Xie, and Daniel H. Huson. 2015. “Fast and Sensitive Protein Alignment Using DIAMOND.” Nature Methods 12 (1): 59–60. 10.1038/nmeth.3176.

5. Cheng, Haoyu, Gregory T. Concepcion, Xiaowen Feng, Haowen Zhang, and Heng Li. 2021. “Haplotype-Resolved de Novo Assembly Using Phased Assembly Graphs with Hifiasm”. Nature Methods 18 (2): 170–75. 10.1038/s41592-020-01056-5.

6. Conesa, Ana, and Stefan Götz. 2008. “Blast2GO: A Comprehensive Suite for Functional Analysis in Plant Genomics.” International Journal of Plant Genomics 2008:619832. 10.1155/2008/619832.

7. Espelta, Josep Maria, Harold Arias-LeClaire, Marcos Fernández-Martínez, Enrique Doblas-Miranda, Alberto Muñoz, and Raúl Bonal. 2017. “Beyond Predator Satiation: Masting but Also the Effects of Rainfall Stochasticity on Weevils Drive Acorn Predation.” Ecosphere 8 (6): e01836. 10.1002/ecs2.1836.

8. Flynn, Jullien M., Robert Hubley, Clément Goubert, Jeb Rosen, Andrew G. Clark, Cédric Feschotte, and Arian F. Smit. 2020. “RepeatModeler2 for Automated Genomic Discovery of Transposable Element Families.” Proceedings of the National Academy of Sciences 117 (17): 9451–57. 10.1073/pnas.1921046117.

9. Gibson, Lester P. 1969. “Monograph of The Genus *Curculio* in the New World (Coleoptera: Curculionidae) Part I. United States and Canada.” In Monograph of The Genus Curculio in the New World (Coleoptera: Curculionidae) Part I. United States and Canada, 6:241. Entomological Society of America. 10.4182/LDPY3443.6-5.241.

10. Gilbert, Clément, Jean Peccoud, and Richard Cordaux. 2021. “Transposable Elements and the Evolution of Insects.” Annual Review of Entomology 66 (Volume 66, 2021): 355–72. 10.1146/annurev-ento-070720-074650.

11. Greiner, Stephan, Pascal Lehwark, and Ralph Bock. 2019. “OrganellarGenomeDRAW (OGDRAW) Version 1.3.1: Expanded Toolkit for the Graphical Visualization of Organellar Genomes.” Nucleic Acids Research 47 (W1): W59–64. 10.1093/nar/gkz238.

12. Guan, Dengfeng, Shane A. McCarthy, Jonathan Wood, Kerstin Howe, Yadong Wang, and Richard Durbin. 2020. “Identifying and Removing Haplotypic Duplication in Primary Genome Assemblies.” *Bioinformatics (Oxford*, England) 36 (9): 2896–98. 10.1093/bioinformatics/btaa025.

13. Higaki, Morio. 2016. “Prolonged Diapause and Seed Predation by the Acorn Weevil, *Curculio robustus*, in Relation to Masting of the Deciduous Oak *Quercus acutissima*.” Entomologia Experimentalis et Applicata 159 (3): 338–46. 10.1111/eea.12444.

14. Hoff, Katharina J., and Mario Stanke. 2019. “Predicting Genes in Single Genomes with AUGUSTUS.” Current Protocols in Bioinformatics 65 (1): e57. 10.1002/cpbi.57.

15. Huang, Neng, and Heng Li. 2023. “Compleasm: A Faster and More Accurate Reimplementation of BUSCO.” Bioinformatics 39 (10): btad595. 10.1093/bioinformatics/btad595.

16. Kelly, Dave. 1994. “The Evolutionary Ecology of Mast Seeding.” Trends in Ecology & Evolution 9 (12): 465–70. 10.1016/0169-5347(94)90310-7.

17. Laetsch, Dominik R., and Mark L. Blaxter. 2017. “BlobTools: Interrogation of Genome Assemblies.” F1000Research. 10.12688/f1000research.12232.1.

18. Laetsch, Dominik R., Georgios Koutsovoulos, Tim Booth, Jason Stajich, and Sujai Kumar. 2017. “DRL/Blobtools: BlobTools v1.0.1.” Zenodo. 10.5281/zenodo.845347.

19. Li, Heng. 2023. “Protein-to-Genome Alignment with Miniprot.” Bioinformatics 39 (1): btad014. 10.1093/bioinformatics/btad014.

20. Li, Qian-Ya, Xing-Hua Hu, De-Chen Liu, Ao Ouyang, Xin Tong, Yong-Jin Wang, Rong Wang, and Xiao-Yong Chen. 2021. “High Diversity and Strong Variation in Host Specificity of Seed Parasitic Acorn Weevils.” Insect Conservation and Diversity 14 (3): 367–76. 10.1111/icad.12462.

21. Maeto, Kaoru, and Kennichi Ozaki. 2003. “Prolonged Diapause of Specialist Seed-Feeders Makes Predator Satiation Unstable in Masting of *Quercus crispula*.” Oecologia 137 (3): 392–98. 10.1007/s00442-003-1381-6.

22. Matsumura, Yoko, Mohsen Jafarpour, Michał Reut, Bardiya Shams Moattar, Abolfazl Darvizeh, Stanislav N. Gorb, and Hamed Rajabi. 2021. “Excavation Mechanics of the Elongated Female Rostrum of the Acorn Weevil *Curculio glandium* (Coleoptera; Curculionidae).” Applied Physics A 127 (5): 348. 10.1007/s00339-021-04353-8.

23. Menu, Frédéric, and Emmanuel Desouhant. 2002. “Bet-Hedging for Variability in Life Cycle Duration: Bigger and Later-Emerging Chestnut Weevils Have Increased Probability of a Prolonged Diapause.” Oecologia 132 (2): 167–74. 10.1007/s00442-002-0969-6.

24. Nelson, L. A., J. F. Wallman, and M. Dowton. 2007. “Using COI Barcodes to Identify Forensically and Medically Important Blowflies.” Medical and Veterinary Entomology 21 (1): 44–52. 10.1111/j.1365-2915.2007.00664.x.

25. Ojo, James Adebayo, M. Carmen Valero, Weilin Sun, Brad S. Coates, Adebayo Amos Omoloye, and Barry R. Pittendrigh. 2016. “Comparison of Full Mitochondrial Genomes for the Rice Weevil, *Sitophilus oryzae* and the Maize Weevil, *Sitophilus zeamais* (Coleoptera: Curculionidae).” Agri Gene 2 (December):29–37. 10.1016/j.aggene.2016.09.007.

26. Schwarzenberger, Anke, Luxi Chen, and Linda C. Weiss. 2020. “The Expression of Circadian Clock Genes in *Daphnia magna* Diapause.” Scientific Reports 10 (1): 19928. 10.1038/s41598-020-77065-3.

27. Sproul, John S, Lindsey M Barton, and David R Maddison. 2020. “Repetitive DNA Profiles Reveal Evidence of Rapid Genome Evolution and Reflect Species Boundaries in Ground Beetles.” Systematic Biology 69 (6): 1137–48. 10.1093/sysbio/syaa030.

28. Sproul, John S., Scott Hotaling, Jacqueline Heckenhauer, Ashlyn Powell, Dez Marshall, Amanda M. Larracuente, Joanna L. Kelley, Steffen U. Pauls, and Paul B. Frandsen. 2023. “Analyses of 600+ Insect Genomes Reveal Repetitive Element Dynamics and Highlight Biodiversity-Scale Repeat Annotation Challenges.” Genome Research 33 (10): 1708–17. 10.1101/gr.277387.122.

29. Stanke, Mario, Oliver Schöffmann, Burkhard Morgenstern, and Stephan Waack. 2006. “Gene Prediction in Eukaryotes with a Generalized Hidden Markov Model That Uses Hints from External Sources.” BMC Bioinformatics 7 (1): 62. 10.1186/1471-2105-7-62.

30. Sudalaimuthuasari, Naganeeswaran, Biduth Kundu, Khaled M. Hazzouri, and Khaled M. A. Amiri. 2024. “Near-Chromosomal-Level Genome of the Red Palm Weevil (*Rhynchophorus ferrugineus*), a Potential Resource for Genome-Based Pest Control.” Scientific Data 11 (1): 45. 10.1038/s41597-024-02910-3.

31. Sylvester, Terrence, Richard Adams, Wayne B Hunter, Xuankun Li, Bert Rivera-Marchand, Rongrong Shen, Na Ra Shin, and Duane D McKenna. 2024. “The Genome of the Invasive and Broadly Polyphagous Diaprepes Root Weevil, *Diaprepes abbreviatus* (Coleoptera), Reveals an Arsenal of Putative Polysaccharide-Degrading Enzymes.” Journal of Heredity 115 (1): 94–102. 10.1093/jhered/esad064.

32. Tamura, Koichiro, Glen Stecher, and Sudhir Kumar. 2021. “MEGA11: Molecular Evolutionary Genetics Analysis Version 11.” Molecular Biology and Evolution 38 (7): 3022–27. 10.1093/molbev/msab120.

33. Trizna, Mike. 2020. “Assembly_stats 0.1.4.” Zenodo. 10.5281/zenodo.3968775. Uliano-Silva, Marcela, João Gabriel R. N. Ferreira, Ksenia Krasheninnikova, Mark Blaxter,

34. Nova Mieszkowska, Neil Hall, Peter Holland, et al. 2023. “MitoHiFi: A Python Pipeline for Mitochondrial Genome Assembly from PacBio High Fidelity Reads.” BMC Bioinformatics 24 (1): 288. 10.1186/s12859-023-05385-y.

35. Yamada, Hirokazu, and Masa-Toshi Yamamoto. 2011. “Association between Circadian Clock Genes and Diapause Incidence in *Drosophila triauraria*.” PLOS ONE 6 (12): e27493. 10.1371/journal.pone.0027493.

